# Structural Bioinformatics of Four Human Aquaporins and Their Water-Soluble QTY Analogs

**DOI:** 10.64898/2026.06.24.734367

**Authors:** Ethan Xiao, Shuguang Zhang

## Abstract

Human <u>a</u>qua<u>p</u>orins (AQPs) are essential membrane channels, yet their inherent hydrophobicity complicates structural and functional studies. We present the systematic application of the QTY code to human AQPs, integrating it with AlphaFold 3 structure prediction to design and validate that four-representative human AQPs (AQP1, AQP3, AQP4, AQP7) can be converted into water-soluble analogs while maintaining their conformation. This approach features a novel platform for editing challenging membrane proteins. The QTY code was applied to the transmembrane regions of the selected four AQPs. Subsequently, the water-soluble QTY analogs of the four AQPs were predicted using AlphaFold 3. The predicted structures were superposed with CyroEM- or X-ray-determined native structures in PyMOL. Further analyses included root-<u>m</u>ean-*s*quare <u>d</u>eviation (RMSD) calculations, visualization of hydrophobic surface reduction, and inspection of conserved protein-ligand binding ability. After applying the QTY code, sequence changes between native AQPs and their QTY analogs was significant (42.86–48.80%). Nevertheless, their structures superposed well in analyses, with only slight deviations (RMSD < 0.6 Å). In addition, the surface hydrophobicity of all QTY-edited AQPs was significantly reduced. Importantly, molecular contacts between the cholesterol ligand and protein were largely preserved for both native AQP1 and its QTY analog. Finally, all AlphaFold3-predicted structures for AQPs have high confidence values (pLDDT > 90; pTM ∼0.83), supporting the reliability of the predicted structures. The findings demonstrate that membrane protein hydrophobicity can be edited and reduced without compromising fold integrity or functional architecture. Integration of the QTY code with AlphaFold 3 affords a high-throughput platform for designing water-soluble, structurally faithful analogs of challenging membrane proteins. Such a strategy can provide a potent platform for detergent-free biochemical studies and water-soluble analogs for therapeutic monoclonal antibody discoveries, thus advancing research of this pharmacologically important protein family.

## Introduction

Approximately ∼30% of proteins in humans, including aquaporins (AQPs), G-protein-coupled receptors (GPCRs) in living systems, etc., are integral transmembrane proteins, yet they account for almost 60% of known drug targets, displaying their critical role in cellular signaling and pharmacology [1–3]. All membrane proteins have a common property: they are insoluble in an aqueous environment because their surfaces are rich in hydrophobic amino acids, such as leucine (L), isoleucine (I), valine (V), and phenylalanine (F). This makes the research on its structure and function particularly difficult [4–6]. The functional characteristics of a protein depend on its protein structure. Obtaining high-resolution structures of a transmembrane protein via cryo-electron microscopy (CryoEM) or X-ray crystallography as well high accuracy prediction of protein structures becomes essential for structure-based drug design, ligand docking, and antibody development [6–10].

The QTY code rational protein editing provides an alternative strategy. Specifically, the QTY code was first reported by Zhang et al in 2018. [11], aimed to develop a theory and protocol, which enables the design of water-soluble analogs of hydrophobic transmembrane proteins through systematically replacing hydrophobic residues with structurally compatible polar amino acids: leucine (L) → glutamine (Q), isoleucine/valine (I/V) → threonine (T), and phenylalanine (F) → tyrosine (Y) [11]. The hydrophobic surface area can be reduced while the helical packing, backbone topology, or functional site geometry are retained, thereby allowing the production of soluble analogs that retain native fold and activity [11–14]. The experimental studies on QTY-engineered chemokine and cytokine receptors maintain ligand-binding capabilities, confirming that this approach for functional membrane proteins is practical [13, 14].

Aquaporins (AQPs) are a large family of membrane proteins that form channels to let water and small molecules pass through cells [16]. Humans have 13 types (AQP0–AQP12), which are divided into two groups: orthodox aquaporins (AQP0, 1, 2, 4, 5, 6, 8) that mainly transport water, and aquaglyceroporins (AQP3, 7, 9, 10) that also allow glycerol and other small molecules to pass [3]. AQPs play important roles in kidney water balance, brain fluid regulation, skin hydration, and fat cell metabolism. When their function is disrupted, it can lead to conditions such as edema, neuromyelitis optica, skin problems, cancer, obesity, and metabolic syndrome, making them important targets for therapy [16, 17].

AQP1, AQP3, AQP4, and AQP7 were chosen for this study due to their functional diversity, structural complexity, disease relevance, and experimental tractability.

AQP1 is a well-studied water channel found in blood vessel endothelial cells, kidney proximal tubules, and red blood cells. It allows rapid water transport across cell membranes, helps endothelial cell migration, and plays an important role in the formation of new blood vessels [3, 16, 17]. The high-resolution CryoEM structure of AQP1 (PDB: 7UZE, 2.4Å) demonstrated that the cholesterol (CLR) ligand is bound to adjacent monomers, forming a clear “glue” binding pocket. This well-determined AQP1structure can serve as a perfect model and the only currently available experimental structures of various AQPs for assessing how QTY substitutions influence protein structure and ligand-binding properties [16, 18].

AQP3 is a representative aquaglyceroporin that can be found in the skin, the kidney, and other tissues. It transports water and glycerol, involved in skin hydration, barrier function, wound healing, and adipocyte energy metabolism. Dysregulation remains implicated in xerosis, psoriasis, renal disorders, and tumor progression. The tetrameric X-ray structure (PDB: 9QSZ, 2.5 Å) features a widened Ar/R selectivity filter accommodating glycerol, making it good candidate for QTY editing in integral helices in membrane [17, 19].

AQP4 is a water-selective aquaporin that is mainly found in astrocytes in the brain. It plays an important role in maintaining brain water balance, removing excess fluid, and supporting glymphatic flow. AQP4 is the main target of autoantibodies in neuromyelitis optica (NMO), and abnormal AQP4 function has been linked to brain edema, stroke, and neuroinflammation. Its high-resolution X-ray structure (PDB: 3GD8, 1.8Å) provides an excellent model for studying the structural effects of QTY substitutions and for developing soluble antigen analogs [9, 20].

AQP7 is found in fat cells, the pancreas, and the kidneys. It transports glycerol out of cells to support fat breakdown and maintain energy balance. Reduced or impaired AQP7 function has been linked to obesity, insulin resistance, and metabolic syndrome. Its X-ray structure (PDB: 6QZJ, 2.2 Å), which contains glycerol existing outside the layer of transmembrane domains, but not physically bound within the channel, provides a useful negative model for examining how QTY substitutions affect transmembrane helix structure, surface properties, and the protein’s ability to transport its substrate [21, 22].

AQP1, AQP3, AQP4, and AQP7 together represent a diverse set of AQPs that span the following: different functions (e.g., water vs. glycerol transport); structural types (e.g., orthodox vs. aquaglyceroporins); disease relevance, and ease of experimental study. Studying them allows a thorough evaluation of how QTY substitutions affect protein folding, surface properties, oligomer assembly, and overall structural stability, while also testing whether they retain ligand binding under biologically relevant conditions.

We combine the QTY code and AlphaFold3 in this structural bioinformatic study for water-soluble analogs of these AQPs. This is the first systematic evaluation of QTY analogs for multiple AQPs. By comparing protein structures, surface hydrophobicity, and analyzing multiple structural features, we may develop a approach to edit QTY-code soluble membrane analogs that retain functional properties. This strategy can support structural research, biochemical studies, and the development of new therapeutic targets.

## Results and Discussion

### Rationale and Design of the QTY Code for Human Aquaporins

The hydrophobic nature of integral membrane proteins, including aquaporins (AQPs), makes them difficult to study, test, and develop as therapeutic targets. These proteins contain many hydrophobic amino acids, such as leucine (L), isoleucine (I), valine (V), and phenylalanine (F), which help them stay embedded in the cell membrane. However, this property also makes them insoluble in water and often requires detergents for purification and analysis, which can affect protein stability and mask important functional regions.

To address this challenge, we applied the QTY code, a protein editing method based on the structural similarities between these hydrophobic amino acids and certain polar amino acids [11, 12, 23, 24]. By QTY code, hydrophobic amino acids of leucine (L), isoleucine (I) & valine (V), and phenylalanine (F) are replaced respectively by hydrophilic glutamine (Q), threonine (T), and by tyrosine (Y), producing water-soluble protein analogs while preserving their overall structure and function.

The QTY code was applied to four representative human AQPs summarized in Table 1: AQP1, AQP3, AQP4, and AQP7. Although the transmembrane regions underwent substantial sequence changes, with 42.86%–48.80% of the residues replaced (Table 2), the QTY analogs retained physicochemical properties similar to those of the native proteins. The isoelectric point (pI) changed very little (e. g., AQP1: 7.14 to 7.14 and AQP4: 7.59 to 7.56) because the amino acid replacements—glutamine (Q), threonine (T), and tyrosine (Y)—are electrically neutral and do not introduce additional charges. The slightly increased molecular weight (MW) (e. g., AQP1: 28.39 kDa to 28.61 kDa), consistent with the slightly higher atomic mass of the substituting residues of Q, T, and Y (e.g., L: 131.17 Da vs. Q: 146.14 Da), reflects the somewhat larger size of the substituted amino acids. By the QTY code design, substitutions of nonpolar amino acids with chemically similar amino acids but containing more water-friendly hydroxyl (-OH) or amide (-NH₂) groups. These changes result in reduced hydrophobic surface area while maintaining its overall conformation. This structural preservation is supported by the very low RMSD values (0.151–0.551Å) observed for the QTY analogs (Table 2).

**Table 1.** The protein names, UniProt ID, and X-ray/CryoEM structure (Å) with PBD ID.

| Name (UniProt ID) | Structure (Å, PDB ID) | Tissue Expression | Medical Relevance |
| --- | --- | --- | --- |
| AQP1 (P29972) | CryoEM (2.4Å, 7UZE) | Erythrocytes, Kidney (PCT, DTL), Choroid plexus, Microvascular Endothelium | **Colton blood group;** Renal water reabsorption; Edema (brain/ocular); Potential cancer biomarker |
| AQP3 (Q92482) | X-ray (2.6Å, 9QSZ) | Skin (Keratinocytes), Kidney (CD), Epididymis, GI Track, Airways | **GIL blood group;** Skin hydration & Wound healing; Glycerol metabolism link to obesity/diabetes |
| AQP4 (P55087) | X-ray (1.8Å, 3GD8) | Brian (Astrocyte End-feet), Spinal Cord, Retina, Skeletal Muscle | **Neuromyelitis Optica (NMO)** target (anti AQP4 IgG); Brain edema; Megalencephalic leukoencephalopathy |
| AQP7 (O14520) | X-ray (1.9Å, 6QZJ) | Adipose Tissue, Testis, Kidney (PST) | **Lipid Metabolism;** Type 2 Diabetes link; Male infertility (glycerol/Sperm motility) |

Note: The lists detailing tissue expression location, medical relevance, and physiological function provided herein are not exhaustive. Updated findings and results are continually emerging from ongoing studies.

**Table 2.** The characteristics of four native human aquaporins and their QTY analogs.

| **Name** | **RMSD**<br>**(Å)** | **PI** | **MW**<br>**(KDw)** | **Hydrophobicity** | **TM Variation**<br>**(%)** | **Overall**<br>**Variation (%)** |
| --- | --- | --- | --- | --- | --- | --- |
| AQP1<sup>CryoEM</sup> | / | 7.14 | 28.3948 | 0.4896 | / | / |
| AQP1<sup>QTY</sup> | 0.310 | 7.14 | 28.6133 | -0.5249 | 42.86 | 17.91 |
| AQP3<sup>X-ray</sup> | / | 6.74 | 31.5438 | 0.5349 | / | / |
| AQP3<sup>QTY</sup> | 0.551 | 6.73 | 31.8963 | -0.5194 | 47.37 | 18.49 |
| AQP4<sup>X-ray</sup> | / | 7.59 | 34.8297 | 0.4201 | / | / |
| AQP4<sup>QTY</sup> | 0.215 | 7.56 | 35.0988 | -0.6358 | 48.80 | 18.89 |
| AQP7<sup>X-ray</sup> | / | 9.03 | 37.2316 | 0.4018 | / | / |
| AQP7<sup>QTY</sup> | 0.265 | 8.87 | 37.6392 | -0.4712 | 46.49 | 15.50 |

Abbreviations: pI, isoelectric point; MW, molecular weight; TM, transmembrane domain; RMSD, root-mean-square deviation; /, not applicable. Note: The four native human aquaporins (integral membrane proteins) and their corresponding water-soluble QTY analogs are listed in the same order as in Figure 1. RMSD values were calculated from structural superpositions after removal of missing residues (unstructured loops) in the experimentally determined native structures (CryoEM or X-ray) and the corresponding regions in the AlphaFold3-predicted QTY analog structures. For structures determined as tetramers, a single monomer was used for alignment. The QTY code led to significant amino acid substitutions in the transmembrane (TM) domains, ranging from 42.86% to 48.80%. Despite these extensive sequence changes, the overall structural variation, as quantified by RMSD, remained low, ranging from 15.50% to 18.89%.

Protein sequence alignments between the natural AQPs and their QTY versions (Figure 1) show that most amino acid changes occur in the transmembrane (TM) helices, highlighted in yellow. The alignment markers (“|” and “*”) indicate that the TM regions are strongly modified (42.86%–48.80% TM variation, Table 2), while the extracellular and intracellular loop regions stay mostly the same. Because the changes are mainly limited to the TM helices, the overall protein structure and the way subunits interact are less likely to be disrupted.

**Figure 1.**
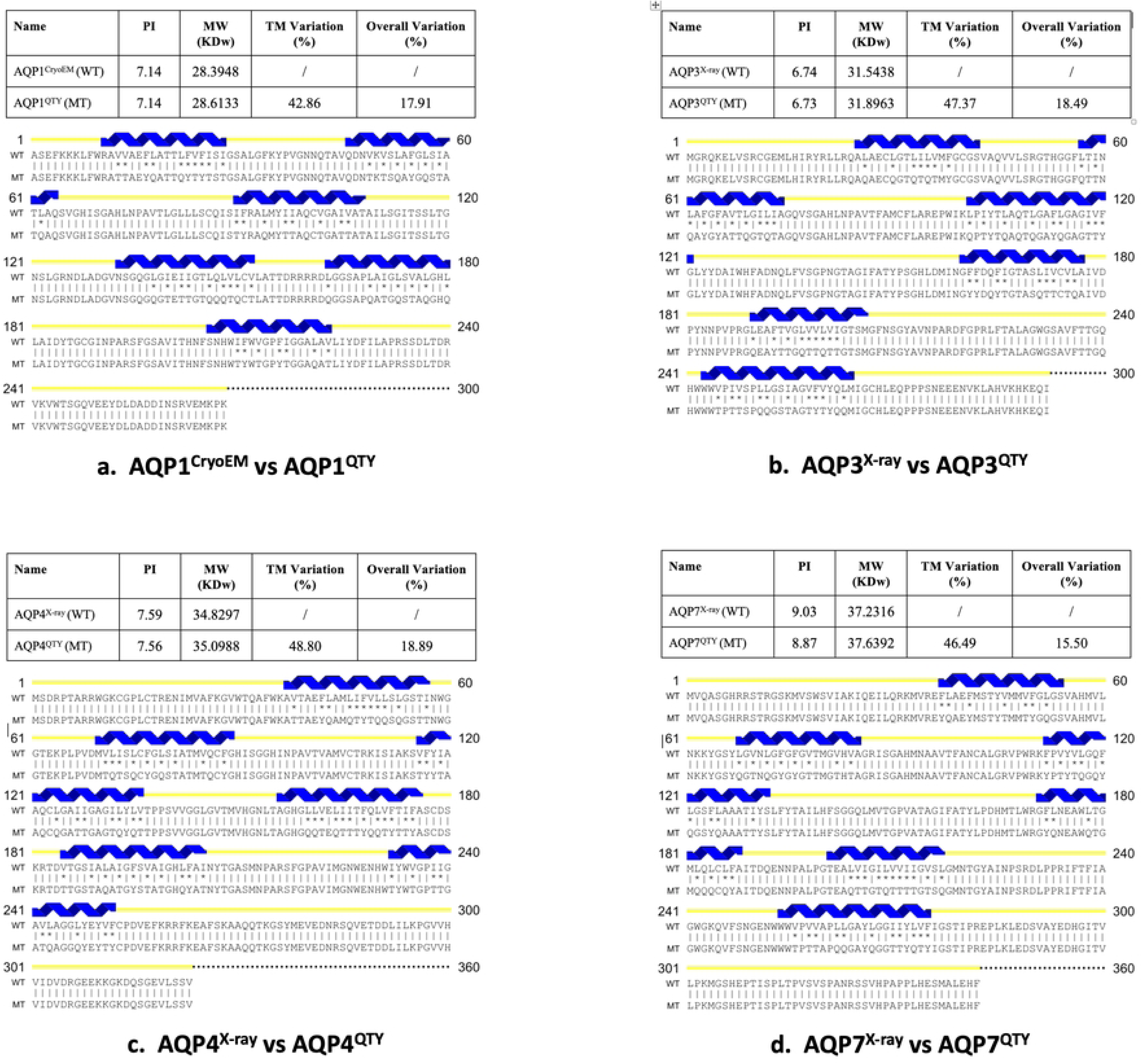
Protein sequence alignments of four native Human Aquaporins with their water-soluble QTY analogs. Protein sequence alignments of native aquaporins and corresponding QTY analogs, designed for reduced hydrophobicity, are shown: (a) AQP1^CryoEM^ vs. AQP1^QTY^; (b) AQP3^X-ray^ vs. AQP3^QTY^; (c) AQP4^X-ray^ vs. AQP4^QTY^; (d) AQP7^X-ray^ vs. AQP7^QTY^. Symbols ‘|’ and ‘*’ within the alignment indicate identical and different amino acids, respectively. QTY substitutions—glutamine (Q), threonine (T), and tyrosine (Y)—replace leucine (L), valine/isoleucine (V/I), and phenylalanine (F) in the native sequences. Predicted transmembrane α-helices are shown in blue above the sequences. Calculated characteristics—including isoelectric point (pI), molecular weight (MW), TM variation %, and overall variation %—are listed above each alignment. Although QTY substitutions introduce substantial changes within TM α-helices (42.86–48.80%), the resulting effects on pI and MW are minimal.

### Validation of AlphaFold3 Prediction Accuracy

AlphaFold3 was used to predict the structures of the QTY analogs, and these predictions were evaluated using several measures to assess prediction accuracy: Predicted Local Distance Difference Test (pLDDT), Predicted Template Modeling (pTM), and Predicted Aligned Error (PAE). The pLDDT measures the confidence in the positioning of each individual residue in the predicted structure [25]. In AlphaFold3 predictions, pLDDT values are color-coded to represent confidence: dark blue regions (pLDDT > 90) indicate very high confidence and are expected to be modeled with high accuracy, light blue regions (pLDDT between 70 and 90) are expected to be modeled well, yellow (pLDDT between 50 and 70) indicates moderate confidence, and orange (pLDDT < 50) suggests low confidence in the accuracy of those regions [9, 10, 26]. pTM evaluates the overall quality of the protein structure by comparing it to known templates [25, 26]. pTM ranges from 0 to 1, in which 1 indicates a perfect structural alignment. This score indicates the strength of topological similarity between experimentally determined protein structures and AlphaFold3 predictions. PAE extends the concept of pLDDT by measuring the predicted distance error between pairs of residues in the structure [26]. This is often displayed as a symmetric matrix, with rows and columns corresponding to residue pairs. Diagonal elements represent the local confidence (predicted errors) for individual residue placement. Darker green regions in the PAE matrix indicate low PAE values, suggesting that AlphaFold3 predicted structure is confident in its relative spatial arrangement.

As visualized in Figure 2, the AlphaFold3 predictions for all four QTY-AQPs exhibited exceptional confidence. Transmembrane regions were predominantly colored dark blue (pLDDT > 90), indicating very high confidence in the atomic positions of these critical structural elements, consistent with the expected behavior for well-structured protein regions. Unstructured loops (loop regions usually display a ribbon-like appearance in the structural model at the ends of the structures, which are intrinsically more flexible) showed lower confidence (yellow/orange, pLDDT < 70), indicating they are intrinsically disordered or unstructured.

**Figure 2.**
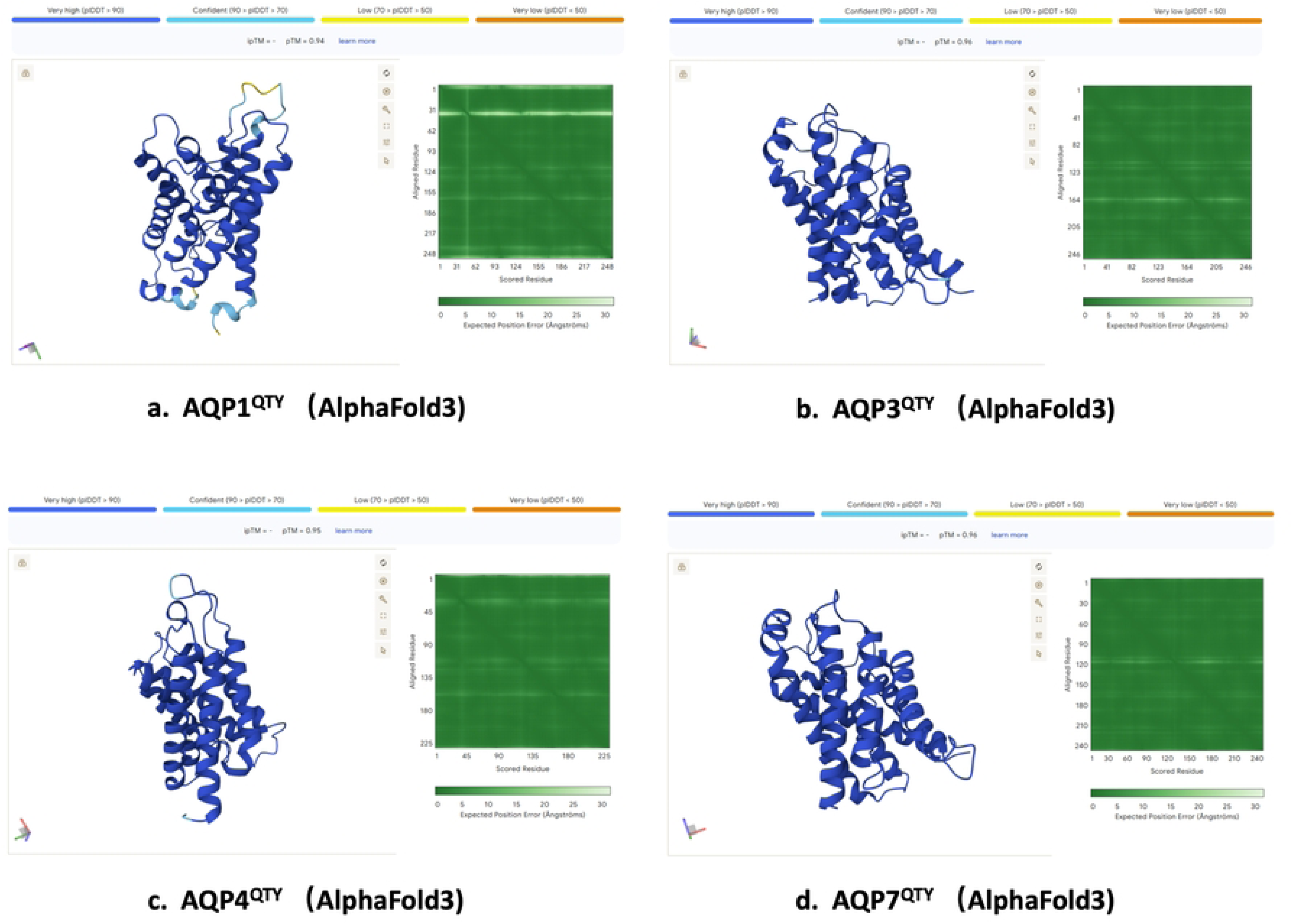
AlphaFold3 prediction quality for water-soluble QTY analogs of four native human aquaporins. Predicted QTY analogs—(a) AQP1^QTY^; (b) AQP3^QTY^; (c) AQP4^QTY^; (d) AQP7^QTY^—are evaluated using AlphaFold3 confidence metrics: per-residue pLDDT, predicted aligned error (PAE) matrices, and overall pTM scores. PAE matrices show predominantly low error (dark green), indicating high confidence in residue pair positions. Transmembrane domains exhibit very high pLDDT (>90, blue/dark blue), while unstructured regions (loops/termini) show lower confidence (<70, yellow/orange). High pTM scores indicate strong structural similarity to known experimental templates.

These unstructured loops were removed in our study for clarity, beginning with the quantitative analysis, to focus on the robust helical core. The overall model quality was further supported by high pTM scores (0.75-0.9, average ∼0.83) as indicated in Figure 2, demonstrating that the models are highly accurate and closely resemble experimentally determined structures.

PAE matrices, as shown in Figure 2, the AlphaFold3 predictions for all four QTY-AQPs displayed large dark green regions, indicating low predicted errors between residue pairs, and confirming high confidence in the relative spatial arrangement of the entire structure. These metrics collectively validate that AlphaFold3 provides highly accurate models suitable for assessing the structural integrity of the QTY-engineered AQPs.

### Structural Superposition of Native and QTY Analog Structures

A primary question of this study was whether the QTY substitutions would preserve the native tertiary and quaternary structures of the aquaporins. We first performed rigorous structural superpositions comparing the AlphaFold3-predicted QTY analogs versus high-resolution experimental structures determined by CryoEM or X-ray crystallography (AQP1: 7UZE; AQP3: 9QSZ; AQP4: 3GD8; AQP7: 6QZJ).

Figure 3 illustrates the near-perfect overlap between the native (magenta) and QTY-analog (cyan) structures. Specific superpositions are as follows: (a) AQP1CryoEM vs AQP1QTY; (b) AQP3X-ray vs AQP3QTY; (c) AQP4X-ray vs AQP4QTY; (d) AQP7X-ray vs AQP7QTY. Analysis of the QTY analogs showed very low RMSD values, all under 0.6 Å: AQP1 (0.310 Å), AQP3 (0.551 Å), AQP4 (0.215 Å), and AQP7 (0.265 Å) (Table 2, Figure 3). Even with many amino acid changes in the transmembrane regions, the structures were only slightly different from those of the native proteins; some key features, such as the six-helix bundle, NPA motifs, and tetrameric assembly, remained intact despite the sequence changes. This shows that AlphaFold3 can accurately predict the structures of QTY analogs. Figure 3 provides evidence that the QTY code can generate water-soluble AQPs without major structural changes.

**Figure 3.**
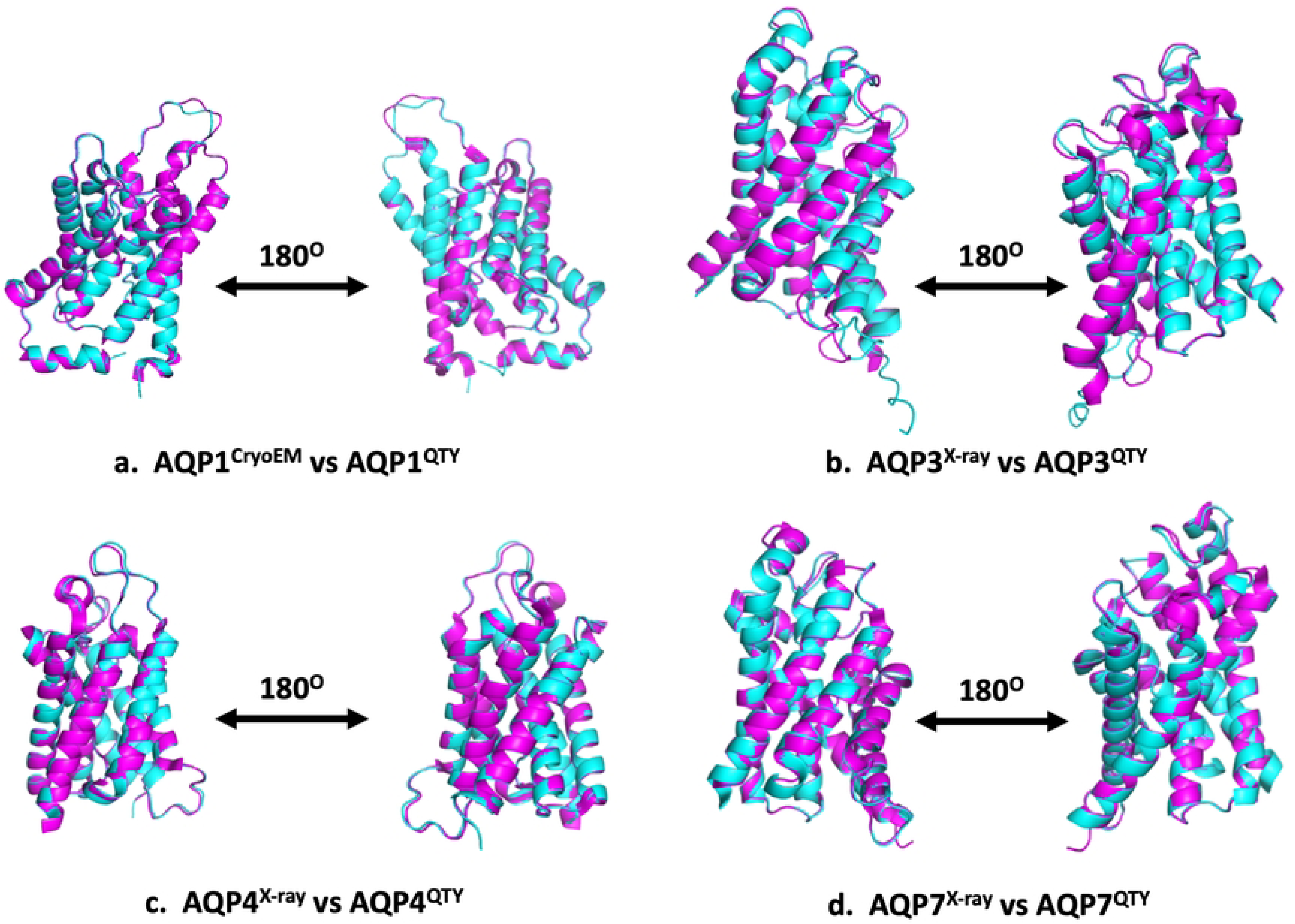
Superposition of four native human aquaporins with their AlphaFold3-predicted water-soluble QTY analogs. Native structures (Cryo-EM/X-ray, magenta) were superimposed with AlphaFold3-predicted QTY analogs (cyan) designed to reduce hydrophobicity. The high degree of structural similarity demonstrates that QTY analogs maintain the native fold. Unstructured loops (flexible loops) with lower confidence (pLDDT < 70) in AlphaFold3 models but absent in experimental structures were removed for clarity. Specific superpositions with RMSD as follows: (a) AQP1^CryoEM^ vs AQP1^QTY^ (RMSD = 0.310 Å); (b) AQP3^X-ray^ vs AQP3^QTY^ (RMSD = 0.551 Å); (c) AQP4^X-ray^ vs AQP4^QTY^ (RMSD = 0.215 Å); (d) AQP7^X-ray^ vs AQP7^QTY^ (RMSD = 0.265 Å).

To better assess the structural integrity of the QTY analogs, we also performed superpositions between the AlphaFold3-predicted native structures (green) and the AlphaFold3-predicted QTY analogs (cyan). Specific superpositions are as follows: (a) AQP1^AlphaFold3^ vs AQP1^QTY^; (b) AQP3^AlphaFold3^ vs AQP3^QTY^; (c) AQP4^AlphaFold3^ vs AQP4^QTY^; (d) AQP7^AlphaFold3^ vs AQP7^QTY^ (Figure 4). Obviously, the quantitative RMSD values remain extremely low (0.150–0.271 Å): AQP1 (0.192 Å), AQP3 (0.271 Å), AQP4 (0.169 Å), and AQP7 (0.150 Å) show exceptional structural overlap, confirming once again that the QTY substitutions do not induce significant conformational changes, and the water-soluble QTY analogs share very similar structures with the native transmembrane AQPs.

**Figure 4.**
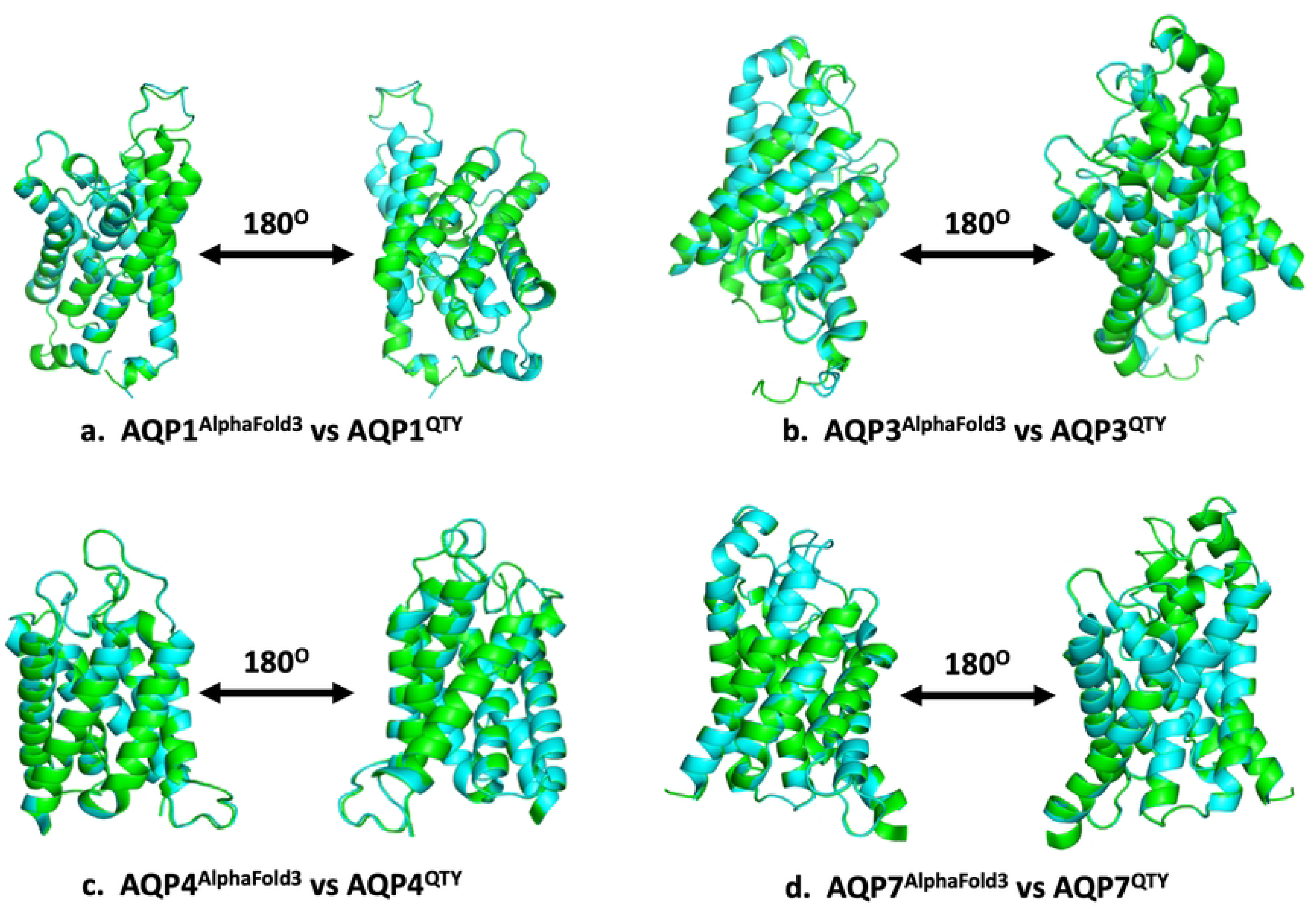
Superposition of AlphaFold3-predicted structures for four human aquaporins and their water-soluble QTY analogs. Native AlphaFold3-predicted structures (green) were superimposed on corresponding QTY analogs (cyan) designed to reduce hydrophobicity. High structural similarity indicates that QTY analogs retain the native fold. Unstructured loops (flexible loops) with lower confidence (pLDDT < 70) in AlphaFold3 models but absent in experimental structures were removed for clarity. The specific superpositions with RMSD as follows: (a) AQP1^AlphaFold3^ vs. AQP1^QTY^, (RMSD = 0.192 Å); (b) AQP3^AlphaFold3^ vs. AQP3^QTY^, (RMSD = 0.271 Å); (c) AQP4^AlphaFold3^ vs. AQP4^QTY^, (RMSD = 0.169 Å); (d) AQP7^AlphaFold3^ vs. AQP7^QTY^, (RMSD = 0.150 Å).

Figure 5 provides further evidence by comparing three versions of each aquaporin: the experimental native structure (magenta), the AlphaFold3-predicted native structure (green), and the QTY analog (cyan). The comparisons include AQP1, AQP3, AQP4, and AQP7. Specific superpositions are as follows: (a) AQP1^CryoEM^ vs AQP1^AlphaFold3^ vs AQP1^QTY^; (b) AQP3^X-ray^ vs AQP3^AlphaFold3^ vs AQP3^QTY^; (c) AQP4^X-ray^ vs AQP4^AlphaFold3^ vs AQP4^QTY^; (d) AQP7^X-ray^ vs AQP7^AlphaFold3^ vs AQP7^QTY^. The close overlap of these three structures shows that AlphaFold3 predicts protein structures with high accuracy. More importantly, the QTY analogs retain the overall quaternary structure and important functional motifs of the native proteins. These results suggest that QTY-engineered AQPs can become water-soluble while maintaining their structural integrity. Such soluble proteins could be easier to express and purify and may be useful in drug discovery, including as antigens for the development of therapeutic monoclonal antibodies.

**Figure 5.**
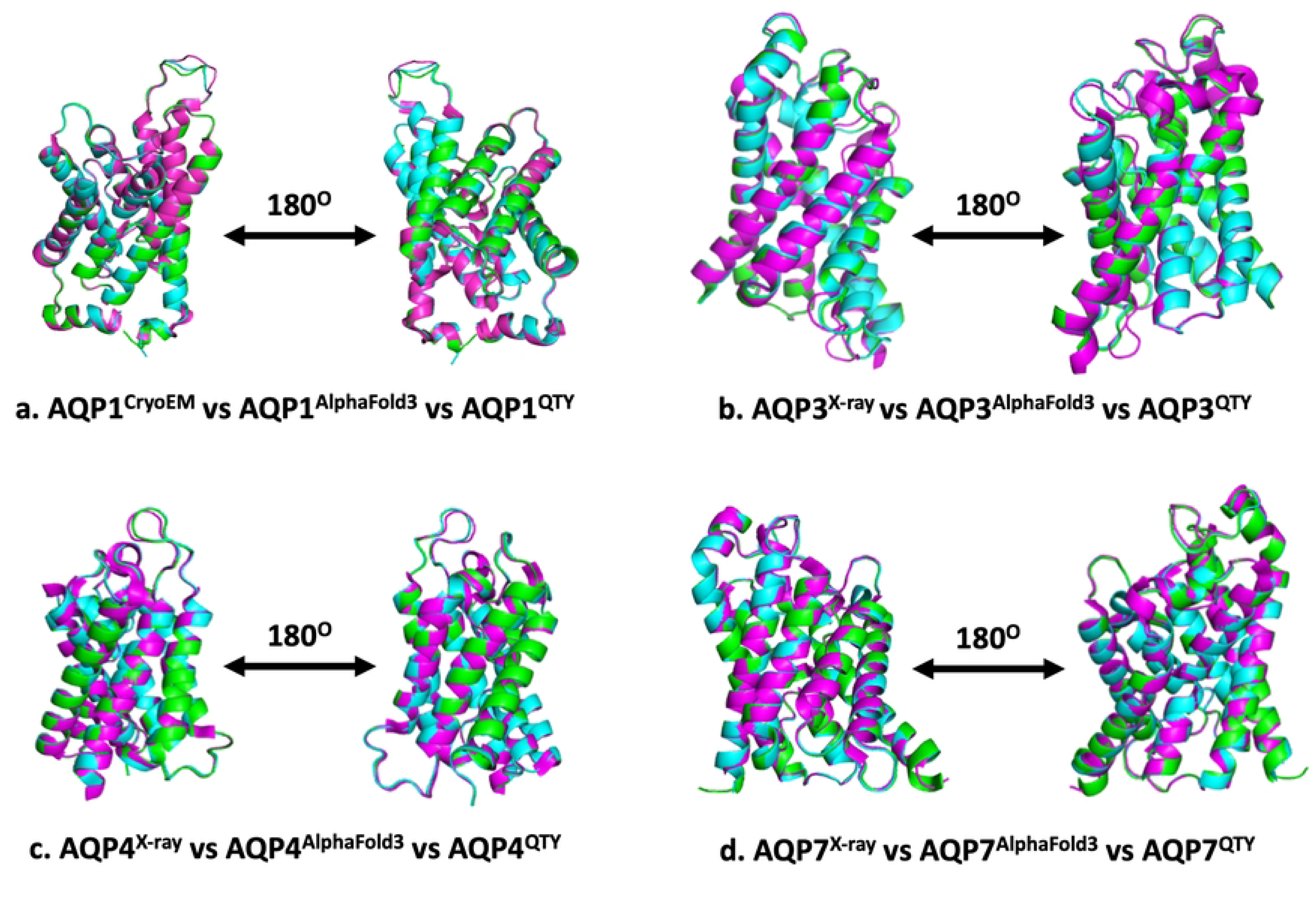
Superposition of four human aquaporins: experimental structures vs. AlphaFold3 predictions and water-soluble QTY analogs. Native structures (Cryo-EM/X-ray, magenta) were compared with AlphaFold3-predicted native structures (green) and QTY analogs (cyan) engineered for reduced hydrophobicity. Unstructured loops (flexible loops) with lower confidence (pLDDT < 70) in AlphaFold3 models but absent in experimental structures were removed for clarity. All three models align closely, showing minimal deviations. These superpositions highlight the accuracy of AlphaFold3 predictions and the structural feasibility of QTY analogs. The specific superpositions as follows: (a) AQP1^CryoEM^ vs. AQP1^AlphaFold3^ vs. AQP1^QTY^; (b) AQP3^X-ray^ vs. AQP3^AlphaFold3^ vs. AQP3^QTY^; (c) AQP4^X-ray^ vs. AQP4^AlphaFold3^ vs. AQP4^QTY^; (d) AQP7^X-ray^ vs. AQP7^AlphaFold3^ vs. AQP7^QTY^.

### Hydrophobic Surface Remodeling

A core objective of the QTY design principle is to convert insoluble, membrane-embedded proteins into soluble, monodisperse forms. This surface transformation is visually demonstrated in Figure 6, which compares the hydrophobicity profiles of three structural models for each target: the experimental native structure, the AlphaFold3-predicted native structure, and the AlphaFold3-predicted QTY analog. The panels show specific comparisons as follows: (a) AQP1^CryoEM^ vs. AQP1^AlphaFold3^ vs. AQP1^QTY^; (b) AQP3^X-ray^ vs. AQP3^AlphaFold3^ vs. AQP3^QTY^; (c) AQP4^X-ray^ vs. AQP4^AlphaFold3^ vs. AQP4^QTY^; (d) AQP7^X-ray^ vs. AQP7^AlphaFold3^ vs. AQP7^QTY^.

**Figure 6.**
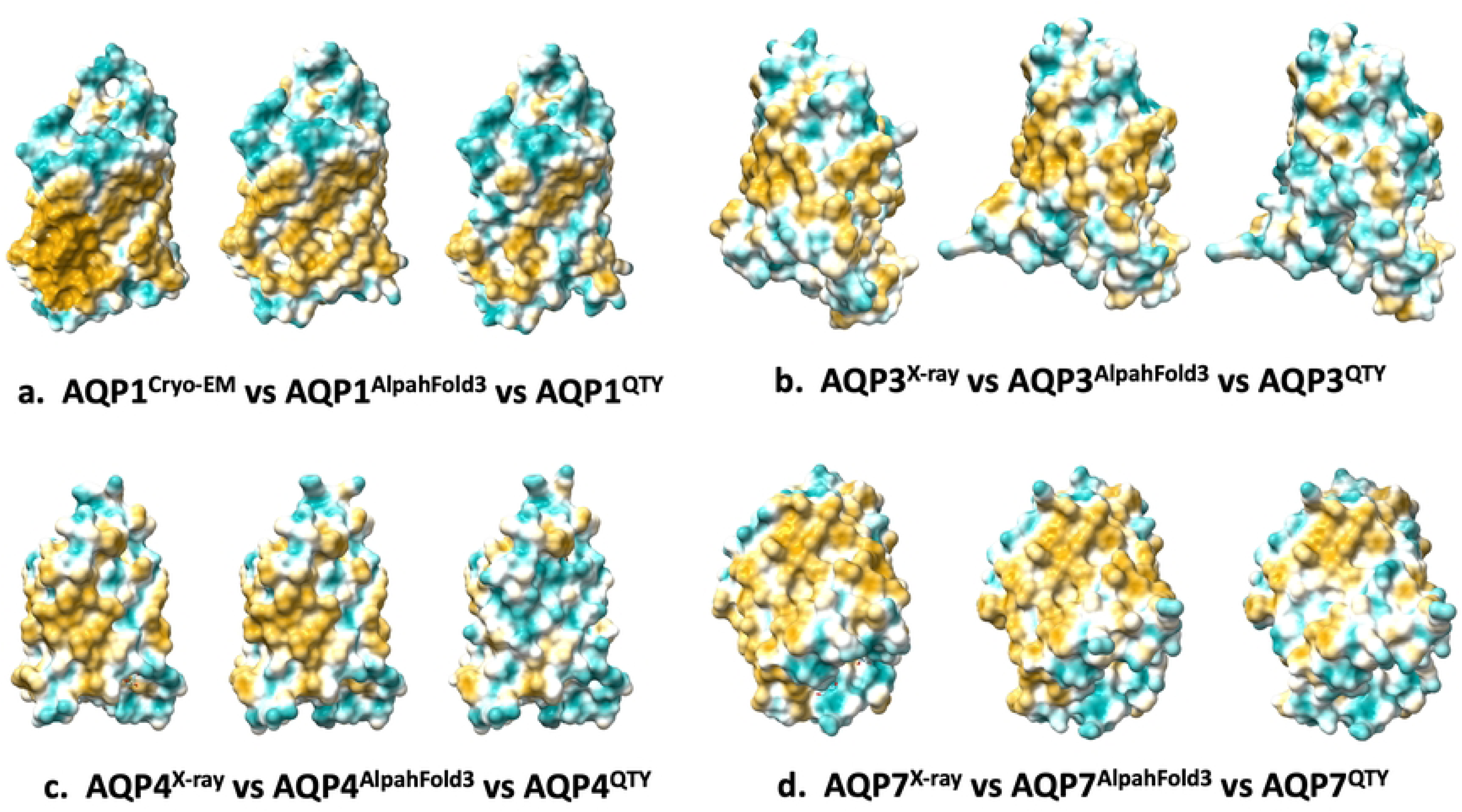
Hydrophobic surface comparison of four human aquaporins: experimental vs. AlphaFold3 predictions and hydrophilic QTY analogs. Native proteins contain hydrophobic residues (L, I, V, F) in the transmembrane helices, shown as yellowish surface patches. Obviously, QTY substitutions (Q, T, Y replacing L, I/V, F) reduce hydrophobicity, making these regions more hydrophilic (cyan). Unstructured loops (flexible loops) with lower confidence (pLDDT < 70) in AlphaFold3 models but absent in experimental structures were removed for clarity. Specific structures for hydrophobicity comparison as follows: (a) AQP1^CryoEM^ vs. AQP1^AlphaFold3^ vs. AQP1^QTY^; (b) AQP3^X-ray^ vs. AQP3^AlphaFold3^ vs. AQP3^QTY^; (c) AQP4^X-ray^ vs. AQP4^AlphaFold3^ vs. AQP4^QTY^; (d) AQP7^X-ray^ vs. AQP7^AlphaFold3^ vs. AQP7^QTY^.

In the 3 structural comparison, both the experimental and AlphaFold3-predicted native structures exhibit extensive yellowish patches across their transmembrane (TM) surfaces. These patches correspond to clusters of hydrophobic, nonpolar residues—including Leucine (L), Isoleucine (I), Valine (V), Phenylalanine (F), Methionine (M), Tryptophan (W), and Alanine (A)—that naturally interact with the lipid bilayer and exclude water.

By applying the QTY code, after replacing key hydrophobic residues (L→Q, I/V→T, F→Y), the QTY analogs look strikingly different with significant reducing yellowish patches. The hydrophobic surface area is reduced, as evidenced by the predominance of cyan surfaces, and diminished yellowish patches. This “hydrophobic-to-hydrophilic” change enables those QTY analogs to dissolve and stay stable in water in the absence of detergents. Importantly, this new water-loving surface doesn’t break the protein’s internal conformation—the core 3D structure remains perfectly intact.

These findings are consistent with previous studies of QTY-engineered chemokine receptors and cytokine receptors, which showed that convert membrane proteins into water-soluble analogs can preserve their structure, stability, and ability to bind their natural ligands [13, 14, 27].

### Preservation of Functional Sites and Ligand-Binding Architecture

To evaluate whether key functional sites of the QTY analogs were retained intact, we used AQP1 as an ideal model due to its experimentally determined and well-characterized cholesterol (CLR) binding pocket at the inter-monomer interface of its tetramer (PDB: 7UZE).

As shown in Figure 7a, the tetrameric structure of native AQP1 determined by CryoEM (PDB: 7UZE) is shown in magenta. It is structurally superimposed on the AlphaFold3-predicted model of the native protein sequence (green) and the AlphaFold 3-predicted model of the engineered, water-soluble QTY analog (cyan). The near-perfect alignment of the three models, with negligible deviations, serves a dual validation. It confirms the high accuracy of the AlphaFold3 prediction for this transmembrane protein and demonstrates that the extensive amino acid substitutions introduced by the QTY code to reduce hydrophobicity do not disrupt the global fold or the quaternary assembly of the protein.

**Figure 7.**
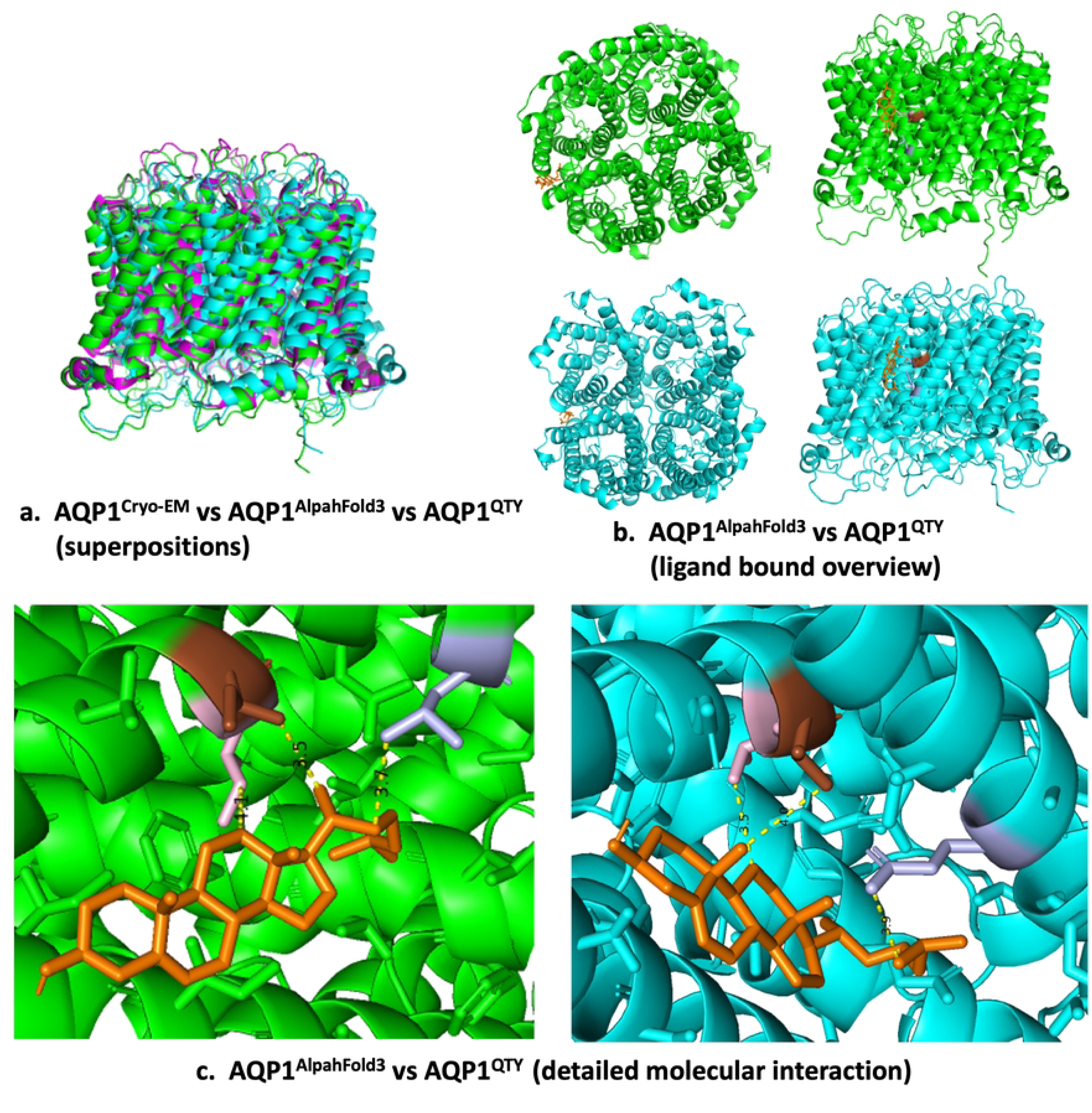
Structural Analysis of Cholesterol Binding in AQP1 and its Water-Soluble QTY Analog. (a) AQP1^CryoEM^ vs. AQP1^AlphaFold3^ vs. AQP1^QTY^ (Superposition): Three tetramers are aligned: the experimentally native structure (magenta), the AlphaFold3-predicted native structure (green), and the water-soluble QTY analog (cyan), showing near-perfect overlap; (b) AQP1^AlphaFold3^ vs. AQP1^QTY^(ligand bound overview): Cholesterol (CLR, orange sticks) sits at the interface between two monomers. The binding position is nearly the same in AlphaFold3-predicted native (green) and the QTY analog (cyan), showing the binding pocket is preserved; (c) AQP1^AlphaFold3^ vs. AQP1^QTY^(detailed molecular interaction): dashed yellow lines mark thee main contacts between cholesterol methyl groups C18, C19, C27, C21 and protein side chains. After using the QTY code (L→Q, I/V→T, F→Y), the original hydrophobic residues (Ile143, Ile144, Leu222 from one monomer, Ile111 from the adjacent monomer) are replaced with polar residues (Thr143, Thr144, Gln222, Thr111). This shows that QTY redesign makes the protein water-soluble without losing its ability to bind cholesterol.

Figure 7b shows a closer view of the cholesterol (CLR) binding site located between two neighboring monomers. The AlphaFold3-predicted native structure (green) and the QTY analog (cyan) are displayed together with the cholesterol molecule (orange). In both structures, cholesterol occupies nearly the same position within the binding pocket, as observed in the experimental 7UZE structure. This suggests that the overall shape and structure of the cholesterol-binding site are maintained in the QTY analog despite the amino acid substitutions.

A close-up, atomic-level view of molecular interactions between the ligand cholesterol (orange sticks) and the AQP1 tetramer is shown for comparison (Figure 7c). The analysis reveals the specific contacts (indicated by yellow dashed lines) that stabilize cholesterol binding in both the AlphaFold3-predicted native model (green) and the QTY analog model (cyan).

In the native structure, the rigid steroid ring of cholesterol forms four important interactions that help stabilize the interface between two neighboring monomers. The C18 methyl group interacts with Ile143 (4.9 Å), the C19 methyl group interacts with Ile144 (7.7 Å), the C27 atom in the cholesterol tail interacts with Leu222 (3.7Å) from one monomer, and the C21 atom interacts with Ile111 (4.8Å) from the adjacent monomer.

In the AQP1 QTY structure, the new, polar amino acids (Thr143, Thr144, Gln222, Thr111) still contact the cholesterol molecule at similar distances (5.7Å, 3.5Å, 3.9Å, and 5.3Å, respectively). Even though the cholesterol shifts slightly, these close contacts demonstrate that the binding “glue pocket” maintains its correct form. Because the new residues can form hydrogen bonds, cholesterol is likely now bound by polar attractions rather than just greasy, hydrophobic forces.

Overall, the ligand-binding study demonstrated that cholesterol binds to the QTY protein in almost the same way. It sits flat, tucked inside a pocket in the protein. Crucially, the network of hydrophobic contacts is preserved in the QTY analog. Even though some parts of the protein were changed from water-repelling (hydrophobic) to water-attracting (polar), the new parts are a similar size and shape. This allows the protein’s pocket to maintain its 3D structure. This demonstrates that the QTY code achieves a “hydrophobic-to-hydrophilic” surface transformation while retaining the structural determinants necessary for specific ligand recognition.

For AQP3, AQP4, and AQP7, there are no experimental ligand-bound structures currently available. However, the conserved NPA motifs and ar/R selectivity filters—the determinants of water and glycerol permeability are clearly preserved in the QTY analogs (Figures 3–5). The structural integrity of these motifs strongly implies that the substrate specificity and transport mechanisms of these aquaglyceroporins are retained in their soluble forms.

### Broader Implications and Applications

This study combines the QTY code with AlphaFold3 to create water-soluble analogs of human aquaporins that closely match their natural structures. AlphaFold3 proves is an accurate protein structure prediction tool, as shown by high pLDDT, pTM, and PAE scores. However, it has limitations, such as possible prediction errors and challenges with modeling ligand interactions. These models are useful for guiding the design of QTY analogs of different membrane proteins [28–30], but laboratory experiments are still needed to confirm the results.

Creating soluble QTY-AQPs solves a major problem in biomedical research. These analogs don’t need detergents, which allows detergent-free biochemical tests, high-resolution structure and other biophysical studies. Because they are soluble and maintain their key surface structures, QTY-AQPs can be used as antigens to help develop therapeutic monoclonal antibodies, such as for neuromyelitis optica (AQP4) or metabolic diseases (AQP7). They also provide a platform for high-throughput drug screening to find new compounds that affect aquaporin function.

The QTY code is a simple and versatile method for turning water-insoluble membrane proteins into soluble, working versions [31–32]. This study creates a trusted computer-aided design process for aquaporins. It opens the door for better 3D structure analysis, function testing, and the creation of new medicines that target these crucial proteins.

For applications, one example is that the QTY code was used to edit CXCR4 water soluble analog. This analog was then recombinantly fused with Fc of IgG and allow self-assembly on 2D lattice of S-layer proteins to create an extremely high-density of CXCR4 analog molecules on top of conducting graphene as electrical sensing device [33]. Such sensing device has extremely high sensitivity to detect natural molecules.

The QTY code can also be reversed as reverse QTY code or rQTY code [34]. We have applied the reverse QTY code to engineer one of the three helical domains of <u>h</u>uman <u>s</u>erum <u>a</u>lbumin (HSA). These edited human serum albumin molecules undergo self-assembly to form well-formed nanoparticles with hydrophobic core. Such nanoparticles has hydrophilic surface and hydrophobic center that can spontaneously encapsulate hydrophobic anti-cancer drugs to form a well-defined complex. Such HSA-drug complex can be delivered into cancer model mice to reduce cancer metastasis [34].

### Summary

In summary, this study successfully extends and validates the QTY code as a generalized strategy for the aquaporin superfamily, building upon its prior success with chemokine and cytokine receptors [13, 14]. By generating hydrophilic, structurally conserved mimics, we provide soluble proteins that hold significant potential as detergent-free reagents for structural and biophysical studies, as ideal antigens for therapeutic monoclonal antibody discovery (e.g., particularly against AQP4 in Neuromyelitis Optica), and as a high-throughput platform for high-throughput drug screening. This work paves a clear way for advanced functional characterization and therapeutic targeting of these critically important but challenging membrane channels.

First, the QTY analogs remained highly similar to their native proteins, with RMSD values below 0.6Å. These results show that important structural features, including the six-transmembrane-helix bundle, tetrameric assembly, and key functional motifs such as the NPA motifs, are preserved even after extensive amino acid substitutions.

Secondly, comparative surface analysis confirmed a dramatic and effective “hydrophobic-to-hydrophilic” transformation, which is the key to achieving aqueous solubility without detergents. Most significantly, detailed investigation of the well-characterized cholesterol-binding site in AQP1 revealed that its architecture remained intact in the QTY analog, indicating preservation of the structural determinants necessary for specific molecular recognition.

Most significantly, detailed investigation of the well-characterized cholesterol-binding site in AQP1 revealed that its architecture remained intact in the QTY analog, indicating preservation of the structural determinants necessary for specific molecular recognition.

Although this computational approach provides strong initial evidence and helps reduce the risks of protein design, experimental validation remains necessary. Future studies should test whether these QTY-AQPs retain their biological functions, especially their ability to transport water and glycerol. The effects of the amino acid substitutions on protein properties should also be investigated. For example, replacing phenylalanine with tyrosine, which has different chemical properties, could affect the behavior of engineered electron-transfer proteins. In addition, further research is needed to determine how the increased hydrophilicity introduced by the QTY code influences cofactor binding, enzymatic activity, and other functions in different membrane protein families.

## Conclusion

Proteins are generally divided into two groups: water-soluble proteins, which are hydrophilic, and membrane proteins, which are hydrophobic. This difference is determined by their amino acid composition and structural properties. In this study, we used the QTY code to convert four hydrophobic human aquaporins—AQP1, AQP3, AQP4, and AQP7—into water-soluble analogs while preserving their overall structures. This work establishes a high-throughput computational approach for designing soluble versions of membrane proteins. Guided by high-confidence AlphaFold3 predictions and validated against experimental structures, our integrated analysis yielded two key findings.

## Method

### The rationale of the QTY code

Purifying and characterizing transmembrane proteins is difficult because they are naturally hydrophobic. As a result, detergents are usually needed to keep them soluble, but this process can be time-consuming and may disrupt their native structure [4]. To overcome this challenge, we used the QTY code, a rational protein engineering method [11]. The QTY code replaces hydrophobic amino acids with structurally similar but more hydrophilic ones: leucine (L) with glutamine (Q), isoleucine (I) and valine (V) with threonine (T), and phenylalanine (F) with tyrosine (Y). Applying these substitutions in the transmembrane regions reduces hydrophobicity while preserving the overall protein structure.

The QTY code was used to create water-soluble versions of four human aquaporins. Even though many amino acids were changed, the QTY analogs retained physicochemical properties very similar to those of the native proteins. As shown in Table 1, the isoelectric point (pI) remained almost unchanged, while the molecular weight (MW) increased only slightly. These results show that the QTY substitutions increase protein hydrophilicity without significantly changing important biophysical properties.

### Protein Sequence and QTY Analog Design

The canonical FASTA sequences for AQP1 (UniProt: P29972), AQP3 (UniProt: Q92482), AQP4 (UniProt: P55087), and AQP7 (UniProt: O14520) were retrieved from the UniProt Knowledgebase (https://www.uniprot.org) [35]. Next, the QTY analogs were designed by substituting all Leu (L) with Gln (Q), Ile (I), and Val (V) with Thr (T), and Phe (F) with Tyr (Y) within the predicted transmembrane helices. These helices were identified using the Protein Solubilizing Server (PSS) (https://pss.sjtu.edu.cn/) [12]. Finally, we computed molecular weights (MWs), isoelectric points (pI), transmembrane (TM) sequence variation, and overall sequence variation using the Expasy Compute pI/MW tool (https://web.expasy.org/compute_pi/) [36].

### AlphaFold3 Structure Prediction

The three-dimensional structures of the four QTY-aquaporin analogs (AQP1^QTY^, AQP3^QTY^, AQP4^QTY^, and AQP7^QTY^) were predicted using the DeepMind AlphaFold3 server (https://alphafoldserver.com) [9]. To validate these predictions, we obtained the corresponding high-resolution experimental structures: AQP1 (PDB: 7UZE, determined by CryoEM), AQP3 (PDB: 9QSZ, determined by X-ray), AQP4 (PDB: 3GD8, determined by X-ray), and AQP7 (PDB: 6QZJ, determined by X-ray) from the RCSB Protein Data Bank (http://www.rcsb.org) [37]. Additionally, predictions for the native protein sequences were sourced from the EBI AlphaFold Protein Structure Database (https://alphafold.ebi.ac.uk) using the corresponding UniProt IDs [10].

### Structural Superposition and Hydrophobicity Analysis

Structural superpositions of the QTY analogs onto their experimental native counterparts were performed in PyMOL v3.0 (https://pymol.org/2) [38]. We used the align command on Cα atoms of the structured core (transmembrane helices and connecting loops, excluding flexible termini). Root-mean-square deviation (RMSD) values were extracted from these alignments.

All structural figures and superpositions were rendered using PyMOL. To visualize and quantitatively analyze hydrophobic surface remodeling, we used UCSF ChimeraX v1.7 (https://www.rbvi.ucsf.edu/chimerax) [39]. Specifically, the molecular surface was colored according to the Kyte-Doolittle hydrophobicity scale [40] using the “Render by Attribute” tool. This allowed us to visually compare the hydrophobic-to-hydrophilic transformation.

### Ligand-Binding Analysis using AutoDock Vina

To test if the QTY engineering aquaporins could still bind their natural ligands, we used computational molecular docking with AutoDock Vina 1.2.3 (https://github.com/ccsb-scripps/AutoDock-Vina) [41]. The analysis focused on AQP1. Its experimental structure (PDB: 7UZE) includes a clearly resolved cholesterol (CLR) molecule in a known binding pocket.

Both the native AQP1 tetramer and the AlphaFold3-predicted AQP1^QTY^ analog were prepared as receptors in PDBQT format using AutoDock Tools (ADT). The cholesterol (CLR) molecule was extracted from the 7UZE structure. Its geometry was optimized, and its rotatable bonds were defined in ADT before saving it in PDBQT format.

The best docking position for native AQP1 was confirmed by comparison with the experimental structure (7UZE). We then examined the top docking positions in the AQP1-QTY analog to determine whether the shape of the binding pocket and its key interactions were preserved.

### Data Availability

A comprehensive suite of publicly accessible bioinformatics resources was utilized in this study. The AlphaFold Protein Structure Database (EMBL-EBI) (https://alphafold.ebi.ac.uk) provided foundational predictions. The structural models (CIF format) for the AQP QTY analogs are available in the GitHub repository https://github.com/Ethan_Xiao/membrane_enzymes, and experimental structures were retrieved from the RCSB Protein Data Bank (http://www.rcsb.org).

For inquiries regarding the QTY-designed models or this study, please contact Ethan Xiao at or Shuguang Zhang at.

## Supplementary Information (SI)

Supplementary data file contains (1) Enlarged alignments of four AQPs with their water-soluble QTY analogs; (2) Membrane topology of four native AQPs; (3) Transmembrane helix predictions of four AQPs with their water-soluble QTY analogs; (4) Enlarged AlphaFold3 predictions confidence metrics (pLDDT, pTM, PAE plots).

## Ethics Approval

Not applicable. This is a computational study that did not involve animal or human subjects.

## Author Contributions

Conceptualization, S.Z.; methodology, E.X., formal analysis, E.X; investigation, E.X; data curation, E.X; writing—review and editing, E.X., S.Z.; visualization, E.X; supervision, S.Z.; project administration, S.Z. All authors have read and agreed to the published version of the manuscript.

